# [^225^Ac]Ac-FAPI-04 Promotes Anti-tumor Immunity via Synergistic Mediated Pattern by *IL-27* Activation and *Tnfaip3* Inhibition

**DOI:** 10.64898/2026.01.07.698054

**Authors:** Shuo Jiang, Siqi Zhang, Tongtong Zhang, Hao Tian, Yekuan Shi, Xin Gao, Suping Li, Rui Wang, Kuan Hu

## Abstract

**Background:** Accumulating evidence substantiates the capacity of targeted radionuclide therapy (TRT) to potentiate tumor immunotherapy responses. However, combining TRT and immunotherapies exhibits substantial varying efficacy, underscoring the importance of investigating mechanisms through which TRT remodels the tumor microenvironment (TME). Despite recent advancements, the specific signaling molecules mediated by TRT that regulate the functions of cytotoxic T lymphocytes (CTLs) remain unclear.

**Methods:** Given the demonstrated potential of α-radionuclides targeting the fibroblast activation protein (FAP) in TRT, this study employed [^225^Ac]Ac-FAPI-04 to investigate the mechanisms of TRT-induced TME remodeling. The cytotoxic effects were first evaluated in the B16F10 cell line, followed by an assessment of its *in vivo* therapeutic efficacy in tumor-bearing mice. RNA-sequencing analysis was utilized to elucidate underlying signaling pathways involving immune responses.

**Results:** [^225^Ac]Ac-FAPI-04 significantly induced DNA double-strand breaks, further elevated ROS levels, and apoptosis in tumor cells. Transcriptomic profiling revealed that [^225^Ac]Ac-FAPI-04 activates the IL-27/JAK-STAT pathway to amplify interferon-mediated responses, indicating IL-27 serves as a critical regulatory signal for CTLs during TRT. A concomitant observation was the downregulation of *Tnfaip3* following [^225^Ac]Ac-FAPI-04 treatment. *Tnfaip3* is implicated in modulating CTLs’ behavior and intratumoral infiltration, while its downregulation was associated with the upregulation of cytotoxic effector molecules, as confirmed in this study.

**Conclusion:** This study identifies a TRT-induced pattern of synergistic signaling mediated by *IL-27* and *Tnfaip3*, which collectively enhances CTL recruitment and potentiates their cytotoxic function. These findings provide a detailed mechanistic rationale for strategies aimed at augmenting the therapeutic efficacy of TRT and immunotherapy combinations.

## BACKGROUND

Targeted radionuclide therapy (TRT) constitutes a promising modality in precision oncology owing to its precise DNA single/double-strand breaks[1-4]. Beyond direct radiation effects, TRT is also recognized for its ability to remodel the tumor microenvironment (TME)[5-7]. For example, sublethal-dose TRT promotes the release of tumor antigens, enhancing immune recognition and triggering systemic anti-tumor immunity[8, 9]. Therefore, the combination of TRT with immune checkpoint blockade (ICB) can increase the therapeutic efficacy of ICB[10, 11]. Besides, some preclinical studies demonstrate that TRT can reshape the TME by upregulating PD-L1 expression and increasing cytotoxic T lymphocytes (CTLs) infiltration[5, 12], thereby achieving complete tumor eradication[13, 14].

However, the intricate crosstalk between immune cells and tumor cells frequently results in heterogeneous efficacy of TRT-ICB combination therapy. In studies of melanoma and breast cancer, combinations of ^225^Ac/^177^Lu/^213^Bi-TRT with ICB showed no significant improvement over monotherapies[15-19]. Some reports even indicate a negative interaction, such as in a breast cancer model combining TRT with anti–PD-1[20]. Although the tumor inhibition efficacy improved following multi-dose TRT combined with anti-PD-1[21, 22], such regimens carry potential risks of bystander effects and drug resistance[23]. Consequently, a deeper mechanistic understanding of how TRT modulates antitumor immunity is essential to reveal critical signals and enhance the efficacy of combined TRT and ICB.

Prior research has shown that TRT activates pathways involved in DNA damage response and immune regulation[6, 24, 25]. Transcriptomic analyses reveal increased NK and CTL infiltration in the TME post-TRT combination with anti-PD-L1. Yet, the precise upstream signals triggered by TRT leading to functional activation of CTLs remain incompletely elucidated. *Interleukin-27* (*IL-27*), a member of the *IL-12*/*IL-6* cytokine superfamily[26, 27], is a critical signal to trigger CTL-mediated immune responses via activating STAT signaling pathways[26, 28].

Considering that antitumor CTLs need *IL-27*, we then hypothesize that *IL-27* is an essential mediator in TRT to elicit CTL-mediated antitumor responses. To validate this, Fibroblast Activation Protein (FAP)-targeted, Actinium-225 (^225^Ac) labeled radioconjugates, [^225^Ac]Ac-FAPI-04[29, 30], were employed as a targeted radionuclide therapy (TAT) agent to investigate the mechanisms underlying TRT-induced remodeling of the TME. This is because TAT characterizes minimizing off-target toxicity and demonstrates remarkable capability in potentiating antitumor immune responses[1]. Firstly, the therapeutic efficacy was confirmed in both B16F10 cells and tumor tissues. Next, RNA-sequencing and molecular experiments analyses demonstrated the critical role of the *IL-27*/*STAT1* axis in TAT-mediated antitumor therapeutics. The enrichment analysis of I*L-27*/*STAT1* pathways also identified *Tnfaip3*, a critical negative regulator of CTLs activation[31], playing a vital role in TAT-associated TME remodeling. The results provide novel insights into how TAT reprograms TEM through *IL-27* activation and *Tnfaip3* suppression, opening a new avenue for improving combination therapies involving IL-27 agonism, ICB, and TRT.

## METHODS

### Radiolabeling and stability of [^22^□Ac]Ac-FAPI-04

^225^Ac was acquired from the Affiliated Hospital of North Sichuan Medical College and dissolved in a 0.2 M ammonium acetate solution. The precursor, FAPI-04, was purchased from Bide Pharmatech Co., Ltd.. The ^225^Ac solution (37 kBq, 0.2 mL), 1.25 M sodium acetate (0.1 mL), 8% sodium ascorbate (0.2 M, 0.1 mL), and 1 mM DOTA-FAPI-04 (0.3 mL) were added and reacted at 90 °C for 30 min. *In vitro* stability was determined by incubating the [^225^Ac]Ac-FAPI-04 (0.005 μCi/mL) in PBS buffer or FBS at 37 °C with maintenance. For *in vivo* stability analysis, [^225^Ac]Ac-FAPI-04 (0.5 μCi) was injected into each mouse intravenously. Followed by incubation for Day 3, the samples were analyzed by radio-TLC.

### Cells and animals

B16F10 cells were cultured in DMEM containing 10% FBS (v/v) at 37 °C in a humidified atmosphere containing 5% CO□. Cells were passaged every 2 days, and fresh medium was changed every day.

All animal procedures were conducted in accordance with guidelines approved by the Animal Care & Welfare Committee of the Institute of Materia Medica, Chinese Academy of Medical Sciences & Peking Union Medical College (IMM-S-25-0104). Animal studies were performed using 6-8-week-old male C57BL6 mice purchased from GemPharmatech Co., Ltd. (Nanjing, China). C57BL6 mice under SPF conditions received subcutaneous injections of 1 × 10^6^ B16F10 cells into the right flank. When tumors reached 150 - 300 mm^3^ (Day 8 - 12 post-inoculation), mice were randomized to treatment groups to minimize size variance[32].

### Half-maximal inhibitory concentration

B16F10 cells were seeded in 96-well plates at a density of 5 × 10^3^ cells per well in complete DMEM supplemented with 10 % FBS and grown for 8-16 h. Cells were then treated with serial dilutions of [^225^Ac]Ac-FAPI-04 and vehicle for 24 h. Cell viability was determined quantitatively via the CCK-8 assay. CCK-8 reagent was supplemented into each well 1 h before the incubation period reached its endpoint. The OD_450_ value in each well was measured using a microplate reader.

### DNA damage assay

B16F10 cells were seeded in 6-well plates and treated with 18.9 μCi/mL [^225^Ac]Ac-FAPI-04 for 24 h. Cells were fixed with 4 % PFA and used as a substrate for a γH2AX DNA damage detection kit after treatment. Briefly, cells were washed 3 times with PBS and permeabilized for 20-30 min in 1 % Triton X-100 at room temperature (RT). Next, B16F10 cells were incubated with rabbit anti-γH2AX antibody overnight at 4 °C in the refrigerator, and incubated with goat anti-rabbit-GFP secondary antibody for 1 h at RT. The cells were subsequently examined using an inverted fluorescent microscope.

### ROS assay

B16F10 cells were seeded at a density of 5 × 10^3^ cells per well in 96-well plates. Upon full cell attachment, the cells were exposed to H_2_O_2_ (120 mM) for 30 min, thereafter, treated with 18.9 μCi/mL [^225^Ac]Ac-FAPI-04, and incubated for 12 hours. DCFH-DA was utilized to assess cellular ROS following the manufacturer’s protocol, incubated in the dark at 37 °C in a cell culture incubator for 20 min, and subsequently washed with PBS 3 times to eliminate excess dye. The cells were subsequently examined using an inverted fluorescent microscope.

### Evaluation of the therapeutic efficacy of [^225^Ac]Ac-FAPI-04

Body weight and tumor dimensions were recorded every 3 days. Tumor volume was measured using digital calipers by recording the length (L) along the longest axis and the perpendicular width (W), calculated by V = 0.5 × L × W^2^. Concurrent body weight was recorded to ± 0.1 g. Humane endpoints were strictly enforced: exclusion for tumor volume > 2,000 mm^3^, ≥ 20% weight loss from baseline, or behavioral distress.

### H&E staining

The B16F10 tumor tissues were collected from tumor-bearing mice after [^225^Ac]Ac-FAPI-04 treatment and fixed with 4% PFA. Deparaffinize and rehydrate formalin-fixed, paraffin-embedded tissue sections through a graded alcohol series to water. The deparaffinized slices were placed in hematoxylin staining solution for 5 min and nucleated and washed with distilled water for 5 min. Fractionation was carried out with 1% ethyl alcohol for several seconds, distilled water soaking for 5 min, thioethyl alcohol 70%, 80% soaking for 5 min, and hematoxylin staining for 5 min. The above steps were repeated, and the stained slices were placed in 95% alcohol, anhydrous ethyl alcohol I, anhydrous ethyl alcohol II, dimethylbenzene I, and dimethylbenzene II for 5 min, respectively, until they were dehydrated and transparent. Seal with medium gum and air dry in the ventilation cabinet. The specimens were observed under a microscope at 200 × magnification, photographed, and analyzed.

### Ki67 immunohistochemical staining

The 4-5 μm sections were deparaffined, rehydrated in a graded series of ethanol and washed in tap water. Antigen retrieval was performed by placing the slides in boiling buffer (0.01 M citrate acid, pH 6) for 15 min. Endogenous peroxidase activity was quenched in methanol with 0.3% H_2_O_2_ for 10 min. Nonspecific binding was blocked at RT for 1 h in PBS-5% BSA. Sections were incubated with an anti-mouse Ki-67 antibody (1:1000 dilution) overnight at 4 °C. Following primary antibody incubation and washing, the slides were treated with an appropriate secondary antibody for 30–60 min at RT. Afterward, the sections were washed 3–5 times with PBS containing Tween 20 (PBST). Visualize with DAB chromogen, which produces a brown precipitate at the antigen site.

### Western blot

B16F10 tumor tissues were lysed with RIPA buffer after [^225^Ac]Ac-FAPI-04 treatment. Followed by centrifuging (12,000 rpm, 20 min) at 4 °C, the supernatant was obtained. The BCA protein assay kit was used to measure the concentration of total protein. The protein samples were electrophoresed on SDS–PAGE with 150 V for 40 min. Then, the protein was transferred to the PVDF membrane at 400 mA for 20 min. The membrane was blocked in 5% skim milk for 2 h at RT and then incubated with primary antibodies (CAS8: 1:1000 dilution, JAK1: 1:1500 dilution, CD8A: 1:1000 dilution, TNFAIP3: 1:1000 dilution, p-STAT1: 1:1000 dilution, GZMB: 1:3000 dilution, FAP: 1:3000 dilution, p-NF-κB: 1:1000 dilution, GAPDH: 1:50000 dilution, β-Actin: 1:50000 dilution) overnight at 4 °C. Then, incubation with rabbit secondary antibodies labeled with HRP (1:10000 dilution) at RT for 1-2 h. The immunoblots were observed by treatment with enhanced chemiluminescence using a Western blotting detection system.

### RNA Extraction and RT-qPCR

Total RNA was extracted from tumor tissues using the RNeasy Plus Mini Kit stored at −80 L. The reverse transcription was subsequently performed using All-in-one 1st Strand cDNA Synthesis Super Mix. The RT-qPCR analysis was then carried out using SYBR on QuantStudio 1. Expression of *Gapdh* was used as a control gene. The RT-qPCR data were processed using the 2^(−ΔΔCT)^ method.**Immunofluorescence staining**

B16F10 tumor tissues were collected from tumor-bearing mice after treatment with informed consent and ethical approval. Frozen samples should be sectioned using a cryostat microtome and kept at RT for 10–15 min, allowing samples to air dry on the slide before fixation in the next step. Cryosections (10 µm) are permeabilized with 0.1% Triton X-100 and blocked with 3% BSA. Primary antibodies are applied for over 10 h at 4°C. After washing with PBS three times, fluorophore-conjugated goat anti-rabbit antibodies are incubated for 1 hour at RT. Slides are stained with DAPI to visualize nuclei and mounted with an anti-fade medium. Imaging is performed using a fluorescence microscope.

### RNA Sequencing and bioinformatics analysis

RNA sequencing was conducted by Shanghai Majorbio Bio-pharm Technology Co., Ltd. Raw reads were aligned to the Mus musculus genome (GRCm38) using HISAT2 v2.2.1, and gene expression quantification was performed using featureCounts v2.0.1.

Differential gene expression analysis was processed using DESeq2 v1.30.1 in R v4.1.0. Genes exhibiting an adjusted *p*-value less than 0.05 and |log□ fold change| greater than 1 were classified as differentially expressed genes. Low-abundance genes (mean counts < 10) were filtered before analysis. Gene Ontology (GO) enrichment used Metascape v3.5 (Mus musculus database) with Biological Process terms filtered by p< 0.01, minimum overlap ≥ 3, and enrichment factor > 1.5; redundant terms were consolidated via semantic similarity clustering (cutoff = 0.7).

Protein-protein interaction networks were generated from the STRING database using differentially expressed genes (DEGs) as input. Interactions with a combined confidence score ≥ 0.7 were retained. Topological analysis included Degree, betweenness, and closeness centrality. Gene Ontology Biological Process terms significantly enriched in DEGs (Metascape output, p < 0.05) were mapped to network nodes. Nodes were colored according to their dominant GO pathway using a discrete color palette.

Relationships between DEGs and enriched GO terms (e.g., Receptor signaling pathway via STAT, Type II interferon production, Keratinocyte differentiation) were illustrated using the GOplot v1.0.2 R package. Gene-term associations were mapped based on statistical significance (*p* < 0.05). Sankey-bubble hybrid plots: Sankey diagrams depicted hierarchical connections between major biological themes, while bubble plots (right panel) quantified enrichment statistics. This composite visualization was generated using ggalluvial v0.12.3 (Sankey) and ggplot2.

Gene Set Enrichment Analysis (GSEA) using clusterProfiler (v4.0.5) was performed on all genes ranked by log□ fold change (descending) against the MSigDB GO Biological Processes subset. Terms with FDR < 0.25 and |NES| > 1.0 were considered enriched.

### Statistical Analysis

Data were analyzed using GraphPad Prism 10.0.2. Comparisons among groups were performed using an unpaired t-test. Data are represented as mean ± SEM. The threshold for statistical significance was set to \**p* < 0.05, \*\**p* < 0.01, \*\*\**p* < 0.001.

## RESULTS

### [^225^Ac]Ac-FAPI-04 induced tumor cell death via DSBs and ROS

Radiolabeling of [^225^Ac]Ac-FAPI-04 was performed, and the products were exposed to PBS and FBS at RT for *in vitro* stability evaluation. The results demonstrated high stability, with over 90% of the compound remaining intact after incubation for 72 h in PBS and FBS (Supplementary Figure S1A-B). Meanwhile, the mouse urine and blood were obtained after 72 h of [^225^Ac]Ac-FAPI-04-injection (Supplementary Figure S1, Supplementary Figure S1C-D), further confirming its satisfactory *in vivo* stability.

We firstly confirmed the expression of FAP in the B16F10 cell line and tumor tissue (Figure 1A). To assess the cytotoxic effects of [^225^Ac]Ac-FAPI-04, B16F10 cells were treated with[^225^Ac]Ac-FAPI-04 in varying concentrations and evaluated 24 h post-treatment (Figure 1B, Supplementary Figure S2). A half-maximal inhibitory effect (IC_50_) was achieved when cells were incubated with 18.97 μCi/mL of [^225^Ac]Ac-FAPI-04 for 24 h. Since TRT is particularly effective in inducing DNA double-strand breaks (DSBs), we examined the expression of DNA damage biomarkers in the [^225^Ac]Ac-FAPI-04-treated cells. Immunofluorescence analysis revealed a significant and time-dependent increase in γH2AX—a specific marker of DSBs—following [^225^Ac]Ac-FAPI-04 treatment (Figure 1C), indicating the induction of clustered DNA damage and sustained activation of the DNA damage response in B16F10 cells, unlike in the control group. Furthermore, we detected reactive oxygen species (ROS) levels in B16F10 cells 24 h after incubation, which showed significantly increased ROS expression levels (Figure 1D), further supporting that the DNA damage occurred in B16F10 cells treated with [^225^Ac]Ac-FAPI-04.

**Fig. 1.**
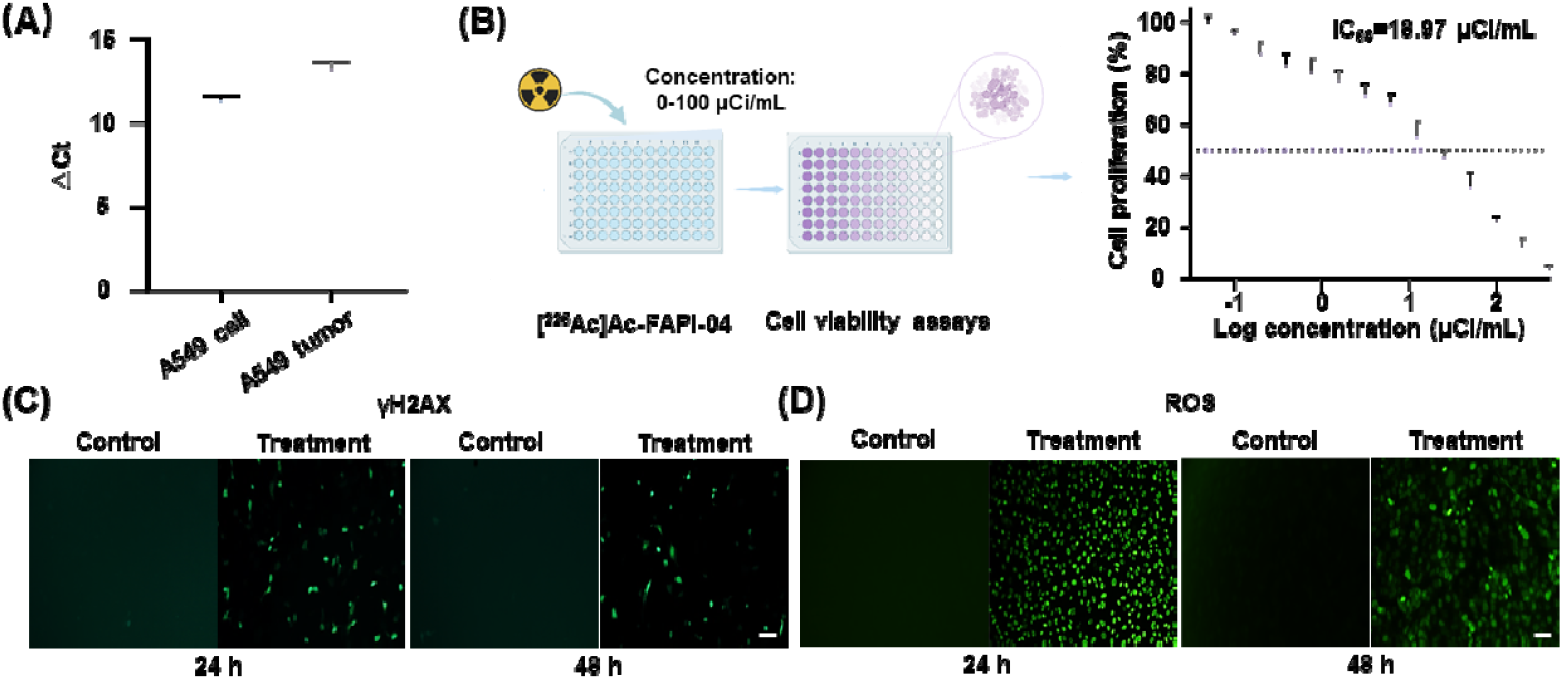
*In vitro* evaluation of cytotoxicity of [^22^□ Ac]Ac-FAPI-04. (A) The gene expression of *Fap* in B16F10 cells and tumor tissue was determined by RT-qPCR. ^Δ^Ct = Ct (*Fap*) − Ct (*Gapdh*). (B) Cytotoxicity analysis of [^225^Ac]Ac-FAPI-04 in B16F10 cells after incubation for 24 h. (C) Fluorescence images of intracellular DNA damage levels (γH2AX) in B16F10 cells following treatment with [^225^Ac]Ac-FAPI-04 or PBS (control) for 24 h and 48 h. Scale bar, 40 μm. (D) Fluorescence images of intracellular ROS levels in B16F10 cells treated with [^225^Ac]Ac-FAPI-04 or PBS (control) for 24 h and 48 h. Scale bar, 40 μm. The error bars indicate the Mean ± SEM (n ≥ 3).

### Therapeutic evaluation of [^225^Ac]Ac-FAPI-04 in B16F10 tumor-bearing mice

The therapeutic efficacy of [^225^Ac]Ac-FAPI-04 was assessed in B16F10 tumor-bearing mice, a model that demonstrates marked FAP expression (Figure 2A-B). Treatment with [^225^Ac]Ac-FAPI-04 resulted in a clear induction of FAP expression compared to the control group, confirming its high targeting specificity and therapeutic potential (Figure 2C). The mice treated with [^225^Ac]Ac-FAPI-04 exhibited significant suppression of tumor growth during 18 days of treatment (Figure 2 D-E). To further evaluate the anti-tumor effect of [^225^Ac]Ac-FAPI-04, tumor tissues were subjected to immunofluorescence and immunohistochemical analysis. The H&E staining indicated that the necrosis of the tumor was significantly increased after [^225^Ac]Ac-FAPI-04 treatment (Figure 2F). Additionally, the positive rate of Ki67 was decreased in the tumor of [^22^□Ac]Ac-FAPI-04 treatment groups (Figure 2G), indicating that [^225^Ac]Ac-FAPI-04 can effectively reduce tumor cell proliferation and induce cell death.

**Fig. 2.**
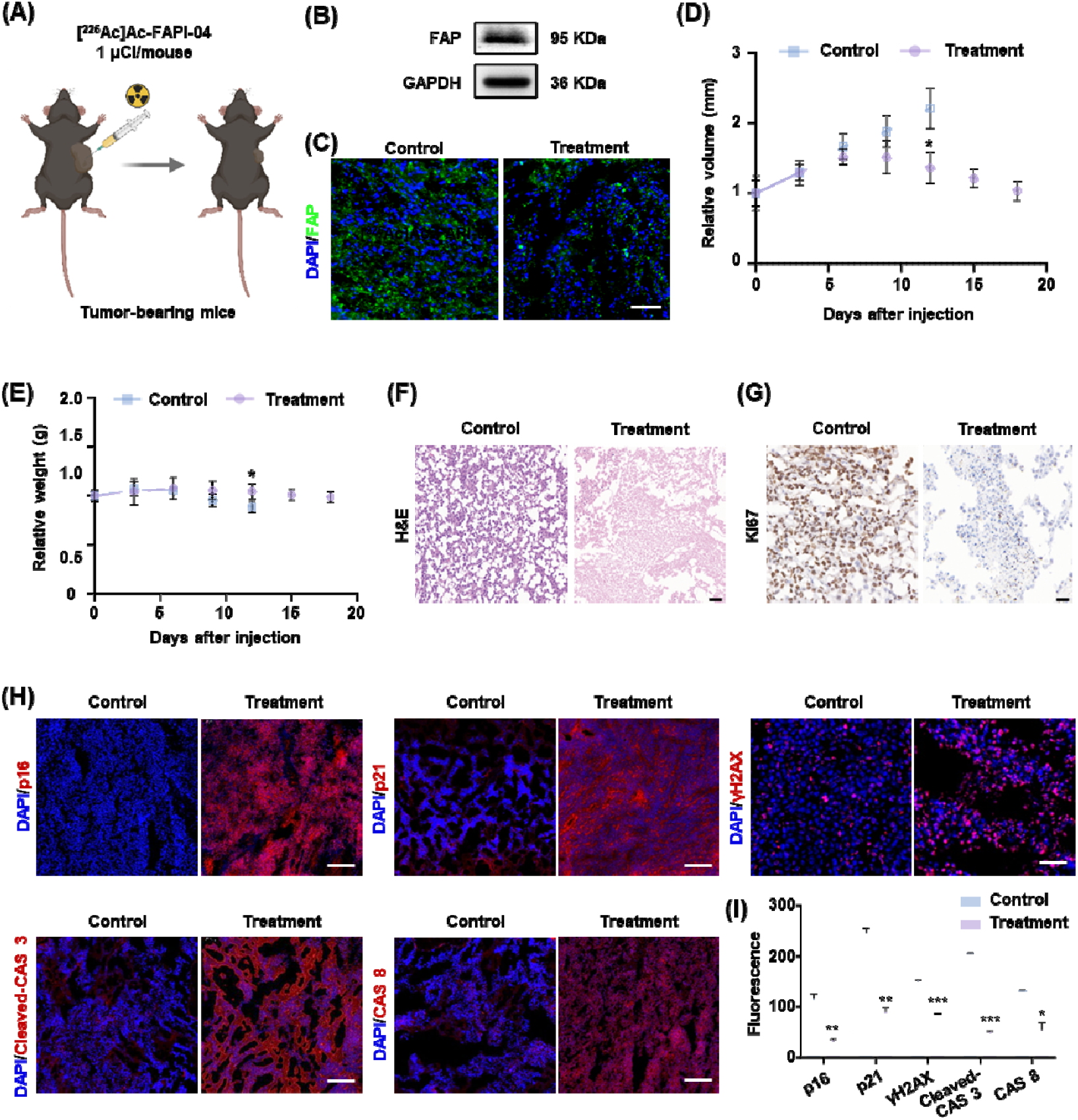
The anti-tumor effect of [^225^Ac]Ac-FAPI-04 in B16F10 tumor-bearing mice. (A) Schematic diagram of the therapy experiment in B16F10 tumor-bearing mice (n ≥ 6). (B) Western blot for FAP in B16F10 tumor tissues before treatment. (C) Immunofluorescence staining for FAP in B16F10 tumor tissues after treatment with [^225^Ac]Ac-FAPI-04 or PBS (control). Scale bar, 100 μm. (D) Relative tumor volume after treatment with [^225^Ac]Ac-FAPI-04 or saline (control). (E) Relative body weight after treatment with [^225^Ac]Ac-FAPI-04 or saline (control). (F) H&E staining of tumor tissue after treatment with [^225^Ac]Ac-FAPI-04 or saline (control). Scale bar, 50 μm. (G) Ki67 staining of tumor tissue after treatment with [^225^Ac]Ac-FAPI-04 or saline (control). Scale bar, 20 μm. (H) Immunofluorescence staining for specific biomarkers of cellular senescence (p16, p21), DNA damage (γH2AX), and apoptosis (cleaved-CAS 3, CAS8). Cleaved-CAS3, cleaved-caspase 3. CAS8, caspase 8. Scale bars, 100 μm. (I) Quantification of mean fluorescence for biomarker expression in Fig. 2(H). The error bars indicate the Mean ± SEM (n ≥ 3). Statistical analysis was conducted using an unpaired t-test. The *p*-value was measured compared with the control group. \**p* <0.05, \*\**p* < 0.01, \*\*\**p* < 0.001.

Consistently, the expression levels of cellular senescence and DNA damage-related biomarkers were significantly increased in tumor tissues following [^225^Ac]Ac-FAPI-04 treatment, including p16, p21, and γH2AX. Concomitantly, the expression of apoptosis-associated biomarkers, such as Cleaved-caspase 3 (Cleaved-CAS3) and Caspase 8 (CAS8), was also markedly upregulated (Figure 2H-I). Taken together, these results confirmed the [^225^Ac]Ac-FAPI-04-induced anti-tumor efficiency.

### Transcriptomic analysis elucidates the anti-tumor effects of ^**225**^**Ac**

To elucidate the therapeutic mechanism of [^225^Ac]Ac-FAPI-04, RNA sequencing was performed on tumor tissues from B16F10 tumor-bearing mice post-treatment with either [^225^Ac]Ac-FAPI-04 or saline (Figure 3A), elucidating the therapeutic mechanism of [^225^Ac]Ac-FAPI-04. A volcano plot of the transcriptome data was generated to identify candidate genes for further functional analysis (Supplementary Figure S3A). Based on these candidates, downstream enrichment analysis was subsequently conducted.

**Fig. 3.**
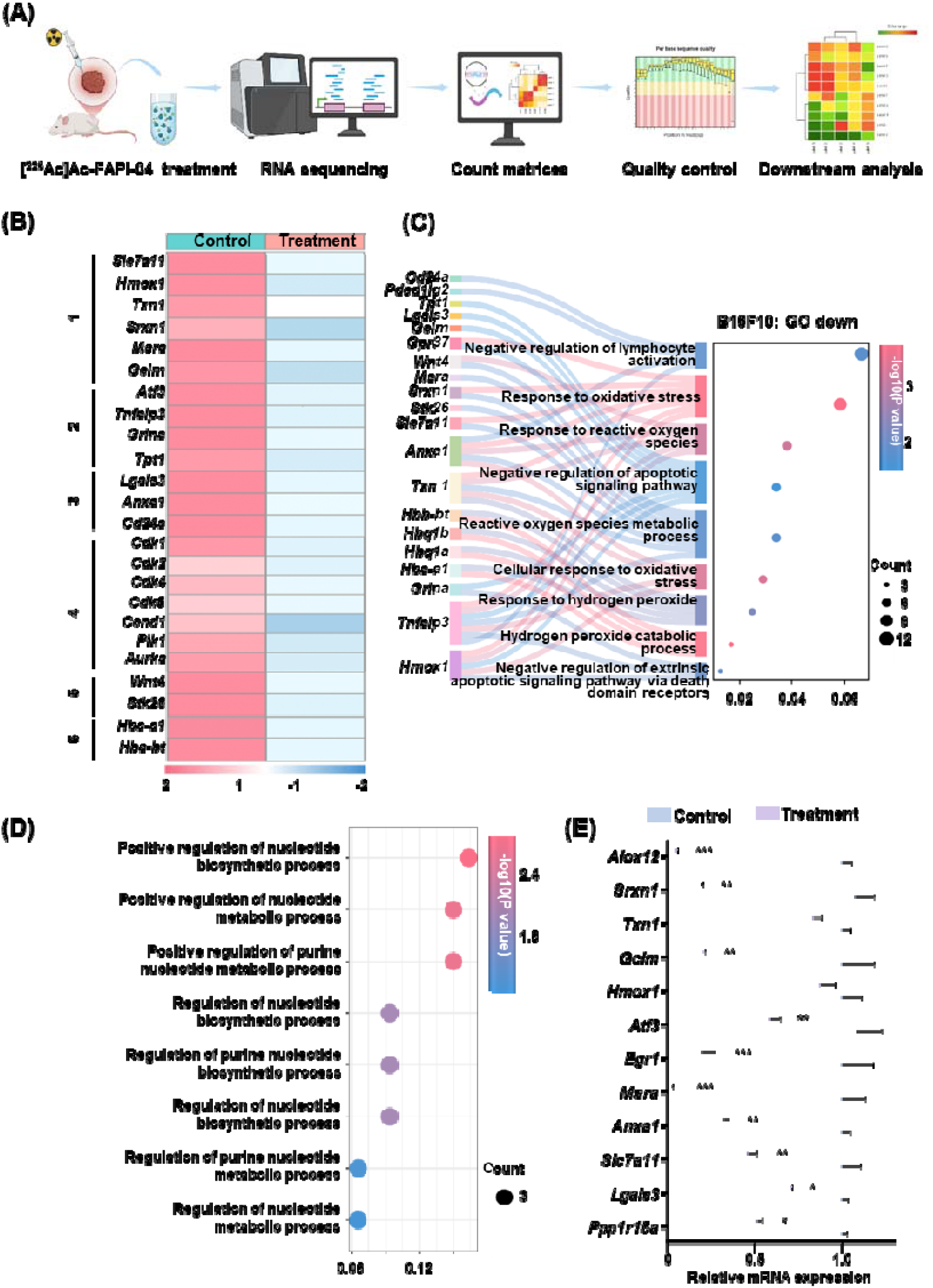
RNA sequencing analysis of genes related to tumor suppression and survival after treatment of [^225^Ac]Ac-FAPI-04. (A) Representative flow chart of the analysis of bulk RNA sequencing (n = 3). (B) Heatmap of the differential expression of common genes related to oxidative stress, signal transduction, immune escape, cell cycle, cell migration, and carcinoembryonic hemoglobin, and values represent the color scale (−2.0 to 2.0). (C) The Sankey diagram showed genes that are downregulated in enrichment GO pathways related to immune response, oxidative stress, and apoptosis. The cutoffs used in the analysis are FDR < 0.05 and q-value < 0.05. (D) GO pathways related to DNA repair. The cutoffs used in the analysis are FDR < 0.05 and |log2 fold change| > 1. (E) Gene expression related to tumor survival determined by RT-qPCR (n = 3). The error bars indicate the Mean ± SEM (n ≥ 3). Statistical analysis was conducted using an unpaired t-test. The *p*-value was measured compared with the control group. \**p* <0.05, \*\**p* < 0.01, \*\*\**p* < 0.001.

To assess the functional impact of [^225^Ac]Ac-FAPI-04 treatment, the enrichment of key DEGs was evaluated across several pathways critically involved in tumor progression, including oxidative stress response, proliferative signal transduction, immune escape, cell cycle control, cell migration, and the regulation of carcinoembryonic hemoglobin. Treatment with [^225^Ac]Ac-FAPI-04 resulted in the significant downregulation of critical genes that sustain intracellular antioxidant capacity, such as *Slc7a11, Hmox1, Txn1, Srxn1, Msra*, and *Gclm* (Figure 3B). Concurrent suppression was observed for pro-tumorigenic genes involved in cell survival and proliferation, including *Atf3, Tnfaip3, Grina*, and *Tpt1* (Figure 3B). Meanwhile, genes that facilitate the immune escape of tumor cells were significantly decreased in expression after treatment (Figure 3B). Furthermore, core cell cycle regulators were also markedly downregulated following treatment (Figure 3B), along with genes related to cell migration and carcinoembryonic hemoglobin production (Figure 3B). Collectively, these findings substantiate the anti-tumor efficacy and immunomodulatory properties of [^225^Ac]Ac-FAPI-04.

Furthermore, GO pathway enrichment analysis of DEGs revealed significant enrichment in biological processes closely associated with tumor inhibition, including immune response, oxidative stress, and apoptosis (Figure 3C, Supplementary Figure S3B). Subsequently, protein-protein interaction analysis of the proteins regulated by these DEGs revealed their enrichment in immune-related pathways (Supplementary Figure S3C-F). These results provide a foundation for further elucidating the mechanism of immune response in the TME. Additionally, significant enrichment was observed in pathways related to nucleotide metabolism and DNA repair, which verified the common killing mechanism of TRT (Figure 3D).

To experimentally validate the RNA-seq findings, the expression of key DEGs was analyzed by RT-qPCR. The results revealed a concordant expression trend with the transcriptomic data (Figure 3E), thereby corroborating the role of [^225^Ac]Ac-FAPI-04 in mediating antitumor efficacy.

### ^225^Ac activates the JAK-STAT pathway by enhancing *IL-27* to regulate CTL cell function

The JAK-STAT signaling pathway is a known contributor to tumor progression and the TME[33, 34]. To investigate the specific role of JAK-STAT in TRT, immunofluorescence staining was performed for CD8 and JAK-STAT pathway components in tumor tissues. Following [^225^Ac]Ac-FAPI-04 treatment, a marked increase was observed in the co-localization of CTLs with JAK1, p-STAT1, and NF-κB (Figure 4A-C), indicating that TRT activated JAK1-STAT1 and NF-κB signaling pathways in CTLs. Moreover, assessment of PD-L1 and p-STAT1 localization in tumor cells revealed increased co-localization following TRT (Figure 4D), implying that activation of the STAT1 pathway might participate in the anti-tumor effect of TRT.

**Fig. 4.**
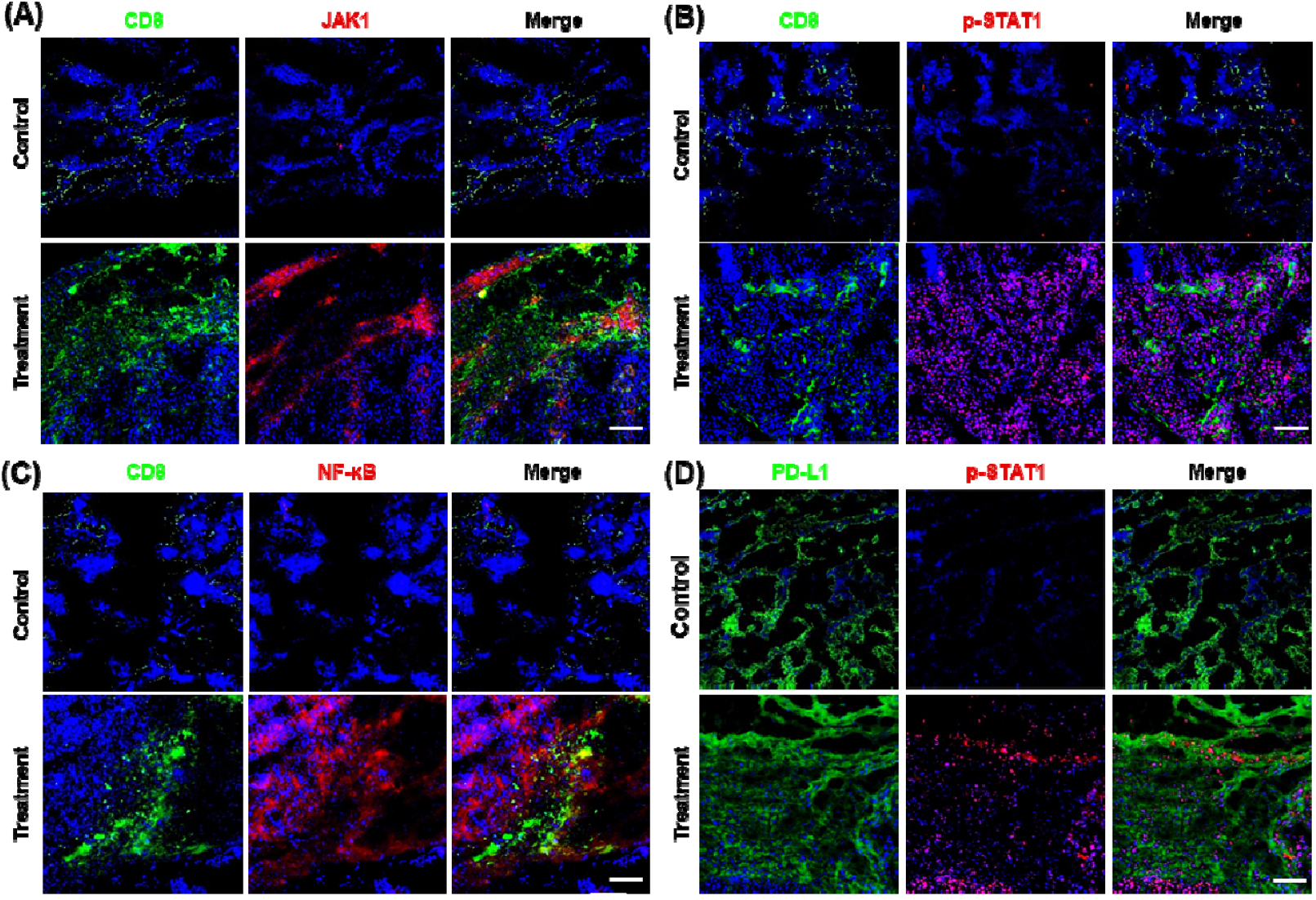
Immunofluorescence staining for co-localization of cellular biomarkers and expression of key proteins in B16F10 tumor tissues after treatment with [^225^Ac]Ac-FAPI-04. (A-C) Co-localization of CD8A with JAK1, STAT1, and NF-κB signaling pathways. (D) Co-localization of PD-L1 with key proteins in STAT1 signaling pathways. Scale bars, 100 μm.

Sequencing analysis identified significant upregulation of JAK-STAT pathway-related genes (e.g. *Stat1, Jak1*) post-treatment with [^225^Ac]Ac-FAPI-04, which was validated by RT-qPCR (Figure 5A-B). Subsequently, enrichment analysis of these genes in GO pathways showed that they were highly enriched in the *IL-27*-related pathway (Figure 5C-D). This is particularly relevant as *IL-27* is known to activate the JAK-STAT pathway, inducing CTLs to secrete granzymes A and B (GZMA/B), thereby enhancing their cytotoxicity and promoting tumor cell killing[35]. Consistent with this reported mechanism, Western blot analysis confirmed markedly elevated protein levels of CD8A, phosphorylated STAT1 (p-STAT1), JAK1, and GZMB after treatment (Figure 5E-F). Collectively, these results confirmed that [^225^Ac]Ac-FAPI-04 can induce the secretion of *IL-27*, then activate the JAK-STAT pathway and enhance the cytotoxic effect on tumor cells.

**Fig. 5.**
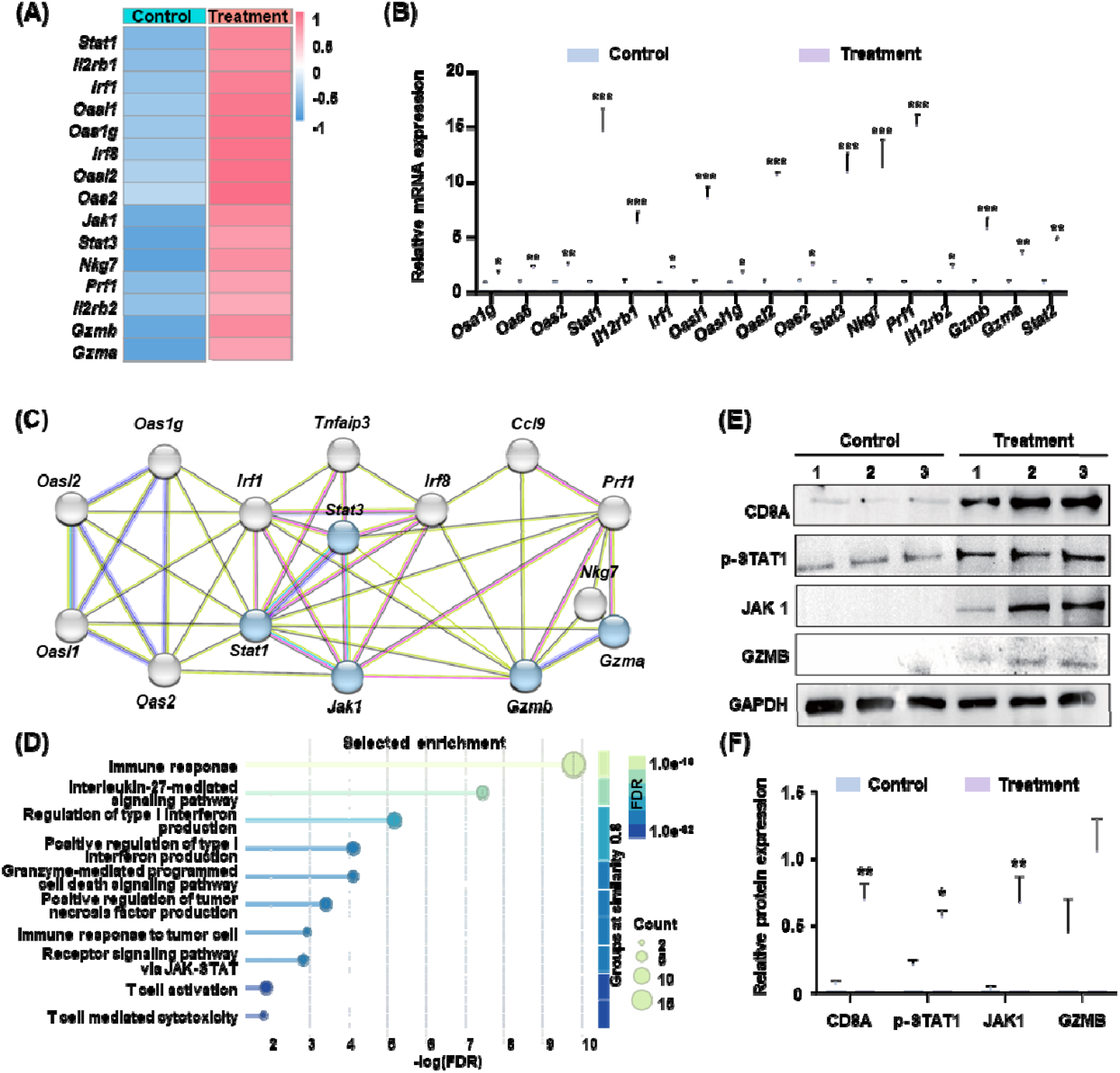
^225^Ac enhances anti-tumor immunity via stimulating IL-27 expression. (A) The heatmap of the expression of genes related to the JAK-STAT pathway, interleukin, and type I interferon (n = 2), and values represent the color scale (−1.0 to 1.0). (B) Gene expression related to the JAK-STAT pathway determined by RT-qPCR (n = 3). The relative expression was measured compared to the control group. (C) The Protein-protein interaction network showed the interactions between proteins related to the JAK-STAT pathway. Confidence score > 0.7 and FDR of GO pathway < 0.05. Created in STRING. (D) The GO enrichment of genes related to the JAK-STAT pathway. The cutoffs used in the analysis are FDR < 0.05 and q-value < 0.05. (E) Expression of CD8A, p-STAT1, JAK1, and GZMB between control and treatment was determined by Western blot. (F) Quantification of protein expression in Fig.5E. Statistical analysis was conducted using an unpaired t-test. The *p*-value was measured compared with the control group. \**p* <0.05, \*\**p* < 0.01, \*\*\**p* < 0.001.

### ^225^Ac enhances anti-tumor immunity by inhibiting *Tnfaip3* and activating the NF-_κ_B signaling pathway

To investigate the mechanisms by which [^225^Ac]Ac-FAPI-04 modulates the TME, GO chord analysis of highly enriched pathways was conducted. This integrated approach identified *Tnfaip3* as the most broadly enriched gene across multiple pathways while exhibiting significant downregulation (Figure 6A). As a crucial negative feedback regulator of TNF signaling, *Tnfaip3* exerts its deubiquitinating enzyme activity to remove critical proteins in the NF-κB signaling pathway, thereby attenuating signal transduction[31].

**Fig. 6.**
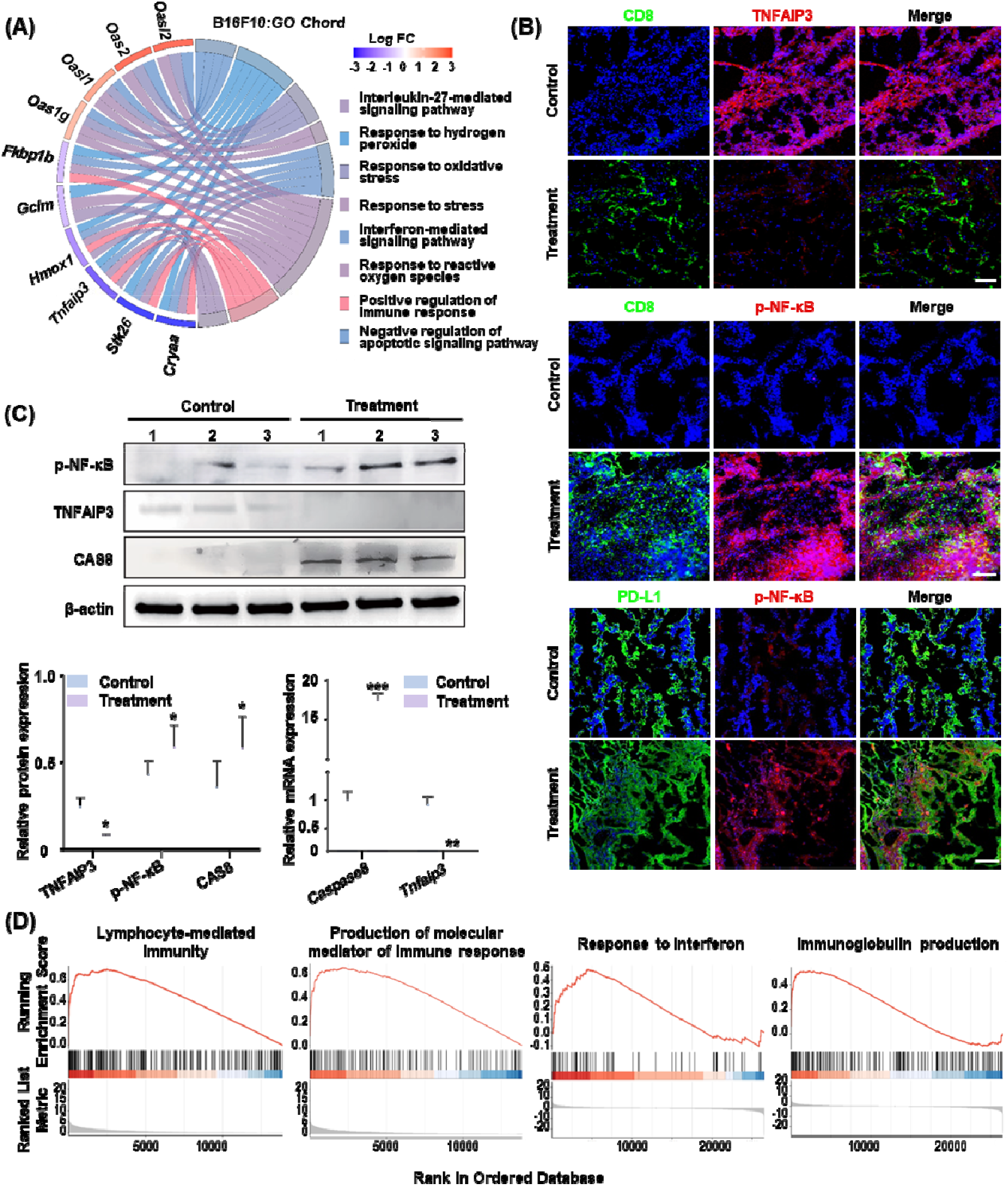
[^225^Ac]Ac-FAPI-04 enhanced anti-tumor immunity via its dual effects on tumor cells and CTLs by TNFAIP3. (A) The GO chord plot of the relationship between upregulated and downregulated genes after [^225^Ac]Ac-FAPI-04 treatment. (B) Co-localization of CD8A with TNFAIP3 or NF-κB and co-localization of PD-L1 with NF-κB. Scale bar, 100 μm. (C) The protein expression of p-NF-κB, TNFAIP3, and CAS8, and the gene expression of *CAS8* and *Tnfaip3* after [^225^Ac]Ac-FAPI-04 treatment (n = 3). (D) GSEA of upregulated and downregulated pathways after [^225^Ac]Ac-FAPI-04 treatment. NES, normalized enrichment score. For each gene, two independent experiments were performed, each with three technical replicates. Statistical analysis was conducted using an unpaired t-test. The *p*-value was measured compared with the control group. \**p* <0.05, \*\**p* < 0.01, \*\*\**p* < 0.001.

The expression level of TNFAIP3 was significantly decreased following treatment with [^225^Ac]Ac-FAPI-04, indicating preserved CTL activation and cytotoxic potential (Figure 6B). There were also strong fluorescent signals of CD8A after treatment with [^225^Ac]Ac-FAPI-04. Furthermore, the partial co-localization of CD8A with p-NF-κB indicated that the NF-κB pathways were specifically activated in CTLs. These imaging demonstrated that CTLs retained their activation and cytotoxic potential following *Tnfaip3* downregulation, which may reflect a productive immune state that could contribute to the inhibition of tumor progression. We revealed significant upregulation of proteins and genes involved in immune regulatory proteins associated with *Tnfaip3* deficiency, including CAS8, phosphate-NF-κB, and CD8A (Figure 6C).

Consistently, GSEA analysis revealed significant activation of pathways such as lymphocyte-mediated immunity and interferon response in the treatment group (Figure 6D), underscoring the role of *Tnfaip3* as a critical node connecting NF-κB and JAK-STAT signaling networks. Thus, TAT may enhance NF-kB signaling and CTLs recruitment by inhibiting *Tnfaip3*, thereby directly eliciting synergic-mediated immune-enhancing effects.

## DISCUSSION

TRT has emerged as a promising strategy for reprogramming the TME from immunosuppressive to immunostimulatory. We confirmed the therapeutic efficacy of [^225^Ac]Ac-FAPI-04, which induced DNA damage, elevated ROS, increased apoptosis, and suppressed tumor growth. Crucially, this study investigated the specific immunomodulatory mechanisms through which TRT regulates CTL function. [^225^Ac]Ac-FAPI-04 showed a key role in activating both the JAK/STAT and NF-κB signaling pathways by enhancing *IL-27* and inhibiting *Tnfaip3*, leading to upregulated interferon-related genes and a potentiated antitumor immune response.

Substantial CTL infiltration was observed post-treatment[13], reinforcing that TRT synergizes with CTL-mediated immunity. RNA-sequencing analysis identified *IL-27* and *Tnfaip3* were key regulators of CTL function in the context of TRT. Treatment with TRT significantly upregulated endogenous *IL-27*, which in turn induced expression of the cytotoxic gene *Gzmb* and enhanced tumor-specific cytotoxic functions—a finding consistent with the regulation pattern involving *IL-27* in anti-tumor immunity[36-38]. We also found that TNFAIP3 expression was markedly downregulated in tumor tissues post-TRT, supporting its role as a genetic determinant of TRT efficacy and the dual regulatory function in both tumor and immune cells[31]. Concurrent suppression of TNFAIP3 in both TRT-exposed tumor cells and CTLs thus appears to act synergistically in enhancing antitumor immunity.

Furthermore, the upregulation of multiple antitumor-related transcription factors and cytokines suggests that [^225^Ac]Ac-FAPI-04 promotes a positive feedback loop and significantly activates anti-tumor responses[39-41]. Taken together, these results demonstrate that *IL-27* and *Tnfaip3*, under the regulation of [^225^Ac]Ac-FAPI-04, function cooperatively to establish a positive feedback circuit between tumor cells and the TME. This interaction initiates a cascade reaction that culminates in the tumor cell. It is necessary to further determine the relationship between the administration duration and dose of [^225^Ac]Ac-FAPI-04 with the immune activation effect, which may potentially change the therapeutic landscape for challenging tumors.

Such insights may ultimately reshape the therapeutic landscape for refractory malignancies. Building upon the central finding that [^225^Ac]Ac-FAPI-04–mediated TRT exerts antitumor effects through the upregulation of *IL-27* and downregulation of *Tnfaip3*, the development of *IL-27* mRNA–based and *Tnfaip3* siRNA–based therapeutics represents a promising strategy to enhance the efficacy of TRT–ICB combination regimens. The administration of exogenous *IL-27* mRNA to tumor tissues may further amplify TRT-induced proliferation of cytotoxic T lymphocytes, offering a viable approach to overcome resistance to immune checkpoint blockade. Similarly, *Tnfaip3* siRNA–based interventions can selectively silence *Tnfaip3* expression in both tumor cells and CTLs, thereby augmenting responsiveness to checkpoint inhibitors. Consequently, *IL-27* mRNA and *Tnfaip3* siRNA agents constitute promising candidates as essential components of multimodal TRT–ICB strategies, with the potential to amplify therapeutic outcomes through a coordinated mechanism of action.

## CONCLUSIONS

The study elucidates the synergistic mechanism by which TAT mediates potent antitumor immunity (Figure 7). We identify *IL-27* and *Tnfaip3* as key signaling mediators responsible for modulating immunoregulatory responses in the context of TRT. Specifically, TAT enhances *IL-27* secretion and triggers interferon-driven cytotoxicity to enhance the anti-tumor immune effect of CTLs. Suppression of *Tnfaip3* by TAT promotes NF-κB phosphorylation, thereby increasing the susceptibility of tumor cells to CTLs. These findings provide a mechanistic basis for enhancing the efficacy of combined TRT-ICB therapy.

**Fig. 7.**
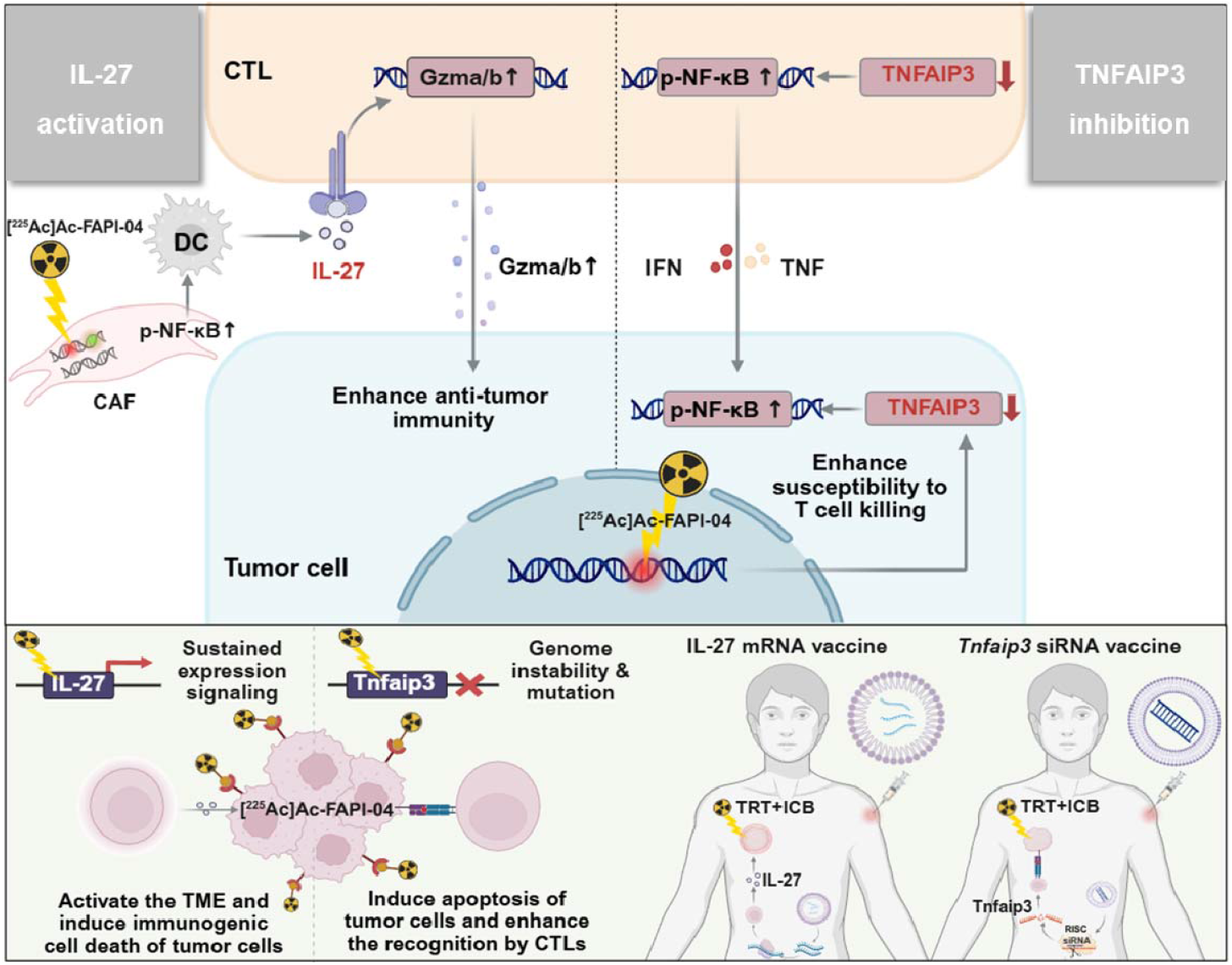
Schematic representation of the anti-tumor immunity activation mechanism of TRT and its translational prospects. The synergic immune activation of [^22^□Ac]Ac-FAPI-04 in CTLs and tumor cells was driven by IL-27 and TNFAIP3. This critical crosstalk directly induced tumor cell death by upregulating the key regulatory molecule *Tnfaip3*, which in turn potentiated the anti-tumor effects of CTLs (above). Future combinations of TRT with immune-modulating strategies (e.g., IL-27 mRNA vaccines, *Tnfaip3* siRNA vaccines) were anticipated to achieve potent anti-tumor effects, advancing personalized tumor therapy (bottom).

## General

We are grateful to all patients who participated in this studyfor their kind cooperation.

## Author contributions

K.H., R.W., and S.Z. contributed to the concept and design of the study. S.J. and H.T. were responsible for acquiring the data. T.Z. and Y.S. performed the data analysis. S.Z., S.J., and T.Z. performed the statistical analysis. R.W. and S.L. contributed to the interpretation of the data. S.J. and T.Z. drafted the manuscript. K.H. and R.W. critically reviewed and approved the manuscript.

## DATA AVAILABILITY

Murine RNA-seq raw data have been deposited into the Sequence Read Archive (SRA) database under the BioProject number PRJNA1347194. Dataverse under the following link: https://dataview.ncbi.nlm.nih.gov/object/PRJNA1347194?reviewer=k5qpkk8917ncpvlmsd0gt2bn1v.

## REFERENCES

1. Zhang S, Wang X, Gao X, Chen X, Li L, Li G, et al. Radiopharmaceuticals and their applications in medicine. Signal Transduction Targeted Ther. 2025;10(1):1.

2. Sun X, Zhang S, Jiang S, Shen J, Wu Y, Zhang H, et al. Combined positron emission tomography-guided modified black phosphorus nanosheet-based photothermal therapy and anti programmed cell death protein ligand 1 therapy. iRADIOLOGY. 2024;2(2):103–12.

3. Zhang S, Ma X, Wu J, Shen J, Shi Y, Wang X, et al. Enhanced radiotheranostic targeting of integrin α5β1 with PEGylation-enabled peptide multidisplay platform (PEGibody): A strategy for prolonged tumor retention with fast blood clearance. Acta Pharm Sini B. 2025;15(2):692–706.

4. Song Z, Zhang J, Qin S, Luan X, Zhang H, Yang M, et al. Targeted alpha therapy: a comprehensive analysis of the biological effects from “local-regional-systemic” dimensions. Eur J Nucl Med Mol Imaging. 2025;Advance online publication.

5. Shi J, Gao H, Wu Y, Luo C, Yang G, Luo Q, et al. Nuclear imaging of PD-L1 expression promotes the synergistic antitumor efficacy of targeted radionuclide therapy and immune checkpoint blockade. Eur J Nucl Med Mol Imaging. 2024;52(3):955–69.

6. Wen X, Shi C, Zeng X, Zhao L, Yao L, Liu Z, et al. A Paradigm of Cancer Immunotherapy Based on 2-[18F]FDG and Anti–PD-L1 mAb Combination to Enhance the Antitumor Effect. Clin Cancer Res. 2022;28(13):2923–37.

7. Zhao L, Pang Y, Zhou Y, Chen J, Fu H, Guo W, et al. Antitumor efficacy and potential mechanism of FAP-targeted radioligand therapy combined with immune checkpoint blockade. Signal Transduction Targeted Ther. 2024;9(1):142.

8. Luri-Rey C, Teijeira Á, Wculek SK, de Andrea C, Herrero C, Lopez-Janeiro A, et al. Cross-priming in cancer immunology and immunotherapy. Nat Rev Cancer. 2025;25(4):249–73.

9. Yang W, Li Y, Gao R, Xiu Z, Sun T. MHC class I dysfunction of glioma stem cells escapes from CTL-mediated immune response via activation of Wnt/β-catenin signaling pathway. Oncogene. 2020;39(5):1098–111.

10. Chen H, Zhao L, Fu K Q L. Integrin αvβ3-targeted radionuclide therapy combined with immune checkpoint blockade immunotherapy synergistically enhances anti-tumor efficacy. Theranostics. 2019;9(25):7948–60.

11. Bao Y, Zhai J, Chen H, Wong CC, Liang C, Ding Y, et al. Targeting m6A reader YTHDF1 augments antitumour immunity and boosts anti-PD-1 efficacy in colorectal cancer. Gut. 2023;72(8):1497–509.

12. Sui H, Guo F, Liu H, Wang R, Li L, Wang J, et al. Safety, pharmacokinetics, and dosimetry of 177Lu-AB-3PRGD2 in patients with advanced integrin αvβ3-positive tumors: A first-in-human study. Acta Pharm Sin B. 2025;15(2):669–80.

13. Zboralski D, Osterkamp F, Christensen E, Bredenbeck A, Schumann A, Hoehne A, et al. Fibroblast activation protein targeted radiotherapy induces an immunogenic tumor microenvironment and enhances the efficacy of PD-1 immune checkpoint inhibition. Eur J Nucl Med Mol Imaging. 2021;50(9):2621–35.

14. Chen J, Zhou Y, Pang Y, Fu K, Luo Q, Sun L, et al. FAP-targeted radioligand therapy with 68Ga/177Lu-DOTA-2P(FAPI)2 enhance immunogenicity and synergize with PD-L1 inhibitors for improved antitumor efficacy. J Immunother Cancer. 2025;13(1):e010212.

15. Chen H, Zhao L, Fu K, Lin Q, Wen X, Jacobson O, et al. Integrin αvβ3-targeted radionuclide therapy combined with immune checkpoint blockade immunotherapy synergistically enhances anti-tumor efficacy. Theranostics. 2019;9(25):7948–60.

16. Vito AA-O, Rathmann S, Mercanti N, El-Sayes N, Mossman KA-O, Valliant J. Combined Radionuclide Therapy and Immunotherapy for Treatment of Triple Negative Breast Cancer. Int J Mol Sci. 2021;22(9):4843.

17. Choi J, Beaino W, Fecek RJ, Fabian KPL, Laymon CM, Kurland BF, et al. Combined VLA-4-Targeted Radionuclide Therapy and Immunotherapy in a Mouse Model of Melanoma. J Nucl Med. 2018;59(12):1843–9.

18. Guzik P, Siwowska K, Fang HY, Cohrs S, Bernhardt P, Schibli R, et al. Promising potential of [177Lu]Lu-DOTA-folate to enhance tumor response to immunotherapy-a preclinical study using a syngeneic breast cancer model. Eur J Nucl Med Mol Imaging. 2021;48(4):984–94.

19. Nosanchuk JD, Jeyakumar A, Ray A, Revskaya E, Jiang Z, Bryan RA, et al. Structure-function analysis and therapeutic efficacy of antibodies to fungal melanin for melanoma radioimmunotherapy. Sci Rep. 2018;8(1):5466.

20. Yang Z, Huang Y, Li F, Hamilton D, Li Z. Small molecule-based angiogenic radionuclide radiation therapy combined with anti-PD1 immune checkpoint blockade for triple-negative breast cancer. J Nucl Med. 2020;61(supplement 1):382.

21. Jiao R, Allen KJH, Malo ME, Rickles D, Dadachova E. Evaluating the Combination of Radioimmunotherapy and Immunotherapy in a Melanoma Mouse Model. J Mol Sci. 2020;21(3):773.

22. Taddio MF, Doshi S, Masri M, Jeanjean P, Hikmat F, Gerlach A, et al. Evaluating [225Ac]Ac-FAPI-46 for the treatment of soft-tissue sarcoma in mice. Eur J Nucl Med Mol Imaging. 2024;51(13):4026–37.

23. Kleinendorst SC, Oosterwijk E, Bussink J, Westdorp H, Konijnenberg MW, Heskamp S. Combining Targeted Radionuclide Therapy and Immune Checkpoint Inhibition for Cancer Treatment. Clinical Cancer Research. 2022;28(17):3652–7.

24. Lejeune P, Cruciani V, Berg-Larsen A, Schlicker A, Mobergslien A, Bartnitzky L, et al. Immunostimulatory effects of targeted thorium-227 conjugates as single agent and in combination with anti-PD-L1 therapy. Journal for ImmunoTherapy of Cancer. 2021;9(10).

25. Zhang J, Yang M, Fan X, Zhu M, Yin Y, Li H, et al. Biomimetic radiosensitizers unlock radiogenetics for local interstitial radiotherapy to activate systematic immune responses and resist tumor metastasis. Journal of Nanobiotechnology. 2022;20(1).

26. Valdés-López JF H-SL, Tamayo-Molina YS, Velilla-Hernández PA, Rodenhuis-Zybert IA, Urcuqui-Inchima S. Interleukin 27, like interferons, activates JAK-STAT signaling and promotes pro-inflammatory and antiviral states that interfere with dengue and chikungunya viruses replication in human macrophages. Front Immunol. 2024;15:1385473.

27. Phan AT, Aunins, E., Cruz-Morales, E., Dwivedi, G., Bunkofske, M., Eberhard, J. N., Aldridge, D. L., Said, H., Banda, O., Tam, Y., Christian, D. A., Vonderheide, R. H., Kedl, R. M., Weissman, D., Alameh, M. G., & Hunter, C. A. The type I IFN-IL-27 axis promotes mRNA vaccine-induced CD8+ T cell responses. bioRxiv. 2025;01(16):633383.

28. Morishima N OT, Asakawa M, Kamiya S, Mizuguchi J, Yoshimoto T. Augmentation of effector CD8+ T cell generation with enhanced granzyme B expression by IL-27. J Immunol. 2005;175(3):1686–93.

29. Piper M, Mueller AC, Karam SD. The interplay between cancer associated fibroblasts and immune cells in the context of radiation therapy. Molecular Carcinogenesis. 2020;59(7):754–65.

30. Allam A, Yakou M, Pang L, Ernst M, Huynh J. Exploiting the STAT3 Nexus in Cancer-Associated Fibroblasts to Improve Cancer Therapy. Frontiers in Immunology. 2021;12.

31. Luo C, Zhang R, Guo R, Wu L, Xue T, He Y, et al. Integrated computational analysis identifies therapeutic targets with dual action in cancer cells and T cells. Immunity. 2025;58(3):745–65.e9.

32. Zhang S, Sun X, Liu W, Wu J, Wu Y, Jiang S, et al. Determining the Multivalent Effects of d-Peptide-Based Radiotracers. Chem Biomed Imaging. 2025;3(3):180–90.

33. Zeng Y, Chen H-q, Zhang Z, Fan J, Li J-z, Zhou S-m, et al. IFI44L as a novel epigenetic silencing tumor suppressor promotes apoptosis through JAK/STAT1 pathway during lung carcinogenesis. Environmental Pollution. 2023;319.

34. Holzgruber J, Martins C, Kulcsar Z, Duplaine A, Rasbach E, Migayron L, et al. Type I interferon signaling induces melanoma cell-intrinsic PD-1 and its inhibition antagonizes immune checkpoint blockade. Nat Commun. 2024;15(1).

35. Bréart B, Williams K, Krimm S, Wong T, Kayser BD, Wang L, et al. IL-27 elicits a cytotoxic CD8+ T cell program to enforce tumour control. Nature. 2025;639(8055):746–53.

36. Koh C-H, Lee S, Kwak M, Kim B-S, Chung Y. CD8 T-cell subsets: heterogeneity, functions, and therapeutic potential. Exp Mol Med. 2023;55(11):2287–2299.

37. Zheng Y, Han L, Chen Z, Li Y, Zhou B, Hu R, et al. PD-L1+CD8+ T cells enrichment in lung cancer exerted regulatory function and tumor-promoting tolerance. iScience. 2022;25(2):103785.

38. Guo F Y. Y. Tumor Necrosis Factor Alpha-Induced Proteins in Malignant Tumors: Progress and Prospects. Onco Targets Ther. 2020;13:3303–3318.

39. Sun Q, Zhao X, Li R, Liu D, Pan B, Xie B, et al. STAT3 regulates CD8+ T cell differentiation and functions in cancer and acute infection. J Exp Med. 2023;220(4):e20220686.

40. Lu C, Klement JD, Ibrahim ML, Xiao W, Redd PS, Nayak-Kapoor A, et al. Type I interferon suppresses tumor growth through activating the STAT3-granzyme B pathway in tumor-infiltrating cytotoxic T lymphocytes. J Immunother Cancer. 2019;7(1):157.

41. Kye Y-C, Lee G-W, Lee S-W, Ju Y-J, Kim H-O, Yun C-H, et al. STAT1 maintains naïve CD8+ T cell quiescence by suppressing the type I IFN-STAT4-mTORC1 signaling axis. Sci Adv. 2021;7(36):eabg8764.

