## Supplement figure for "[^225^Ac]Ac-FAPI-04 Promotes Anti-tumor Immunity via Synergistic Mediated Pattern by *IL-27* Activation and *Tnfaip3* Inhibition"

**Kuan Hu**, Ph.D.

State Key Laboratory of Bioactive Substance and Function of Natural Medicines, Institute of Materia Medica, Chinese Academy of Medical Sciences and Peking Union Medical College, Beijing, China.

**Rui Wang**, Ph.D.

State Key Laboratory of Bioactive Substance and Function of Natural Medicines, Institute of Materia Medica, Chinese Academy of Medical Sciences and Peking Union Medical College, Beijing, China.

**Suping Li,** Dr.

Department of Nuclear Medicine, Affiliated Hospital of North Sichuan Medical College, Nanchong, P.R. China.

**Supplementary material and method**

**Reagent and equipment**

FAPI-04 was purchased from Bide Pharmatech Co., Ltd. (Shanghai, China). DMEM (C11995500BT) was acquired from Thermo Fisher Scientific (Shanghai, China). Fetal bovine serum (FBS, SE100-011) was acquired from VisTech™(AUS). RIPA Lysis Buffer (R0010) and Triton X-100 (T8200) were purchased from Solarbio (Beijing, China). CCK-8 (C0052S), γH2AX DNA damage detection kit (C2035), DCFH-DA (S0034S), and hematoxylin staining solution (C0107) were bought from Beyotime (Shanghai, China). RIPA buffer (RM02998), DAPI (RM02978), anti-Ki67 mouse mAb (A26239), phospho-STAT1 rabbit monoclonal antibody (AP1000), Caspase-8 rabbit monoclonal antibody (A0215), JAK1 rabbit monoclonal antibody (A11963), TNFAIP3 rabbit monoclonal antibody (A2127), CD8A rabbit monoclonal antibody (A23305PMa), FAP rabbit monoclonal antibody (A23789), GAPDH rabbit mAb (A19056), and β-actin rabbit mAb (AC038) were purchased from ABclonal (Wuhan, China). The PD-L1 mouse monoclonal antibody (66248-1-Ig) was bought from Proteintech (Wuhan, China). The phosphate-NF-kB rabbit monoclonal antibody (F0155) was bought from Selleck (USA). The BCA protein assay kit (ZJ102) was purchased from Epizyme (Shanghai, China). The BSA (GC305010) and 4 % PFA (G1101) were brought from Servicebio (Wuhan, China). The RNeasy Plus Mini Kit (DP451) was bought from TIANGEN (Beijing, China). The All-in-one 1st Strand cDNA Synthesis SuperMix (E047-01B) and SYBR (E401-01B) were purchased from Novoprotein (Suzhou, China). The primers were synthesized by Tsingke Biotechnology Co., Ltd. (Beijing, China).

The purification of [²²⁵Ac]Ac-FAPI-04 was analyzed by radio-TLC (BioScan, USA). The OD values of CCK-8 and Ros were detected by a microplate reader from Thermo Fisher (USA). The RT-qPCR was run in QuantStudio 1, and the results were analyzed using QuantStudio software (ThermoFisher, USA). The cell was imaged in an inverted fluorescent microscope (Carl Zeiss, USA). Immunoblots were detected using a Tanon 5200 Western blot imaging system (Tanon). Immunofluorescence staining was imaged using a Nikon A1R SI Confocal from Japan.

**RT-qPCR**

The primers utilized for RT-qPCR are as follows:

| Gene | Forward | Reverse |
| --- | --- | --- |
| *Fap* | GACGGGGGACTGACTTTCTG | TCCCCTCCTTACTTGCGAGA |
| *Tnfaip3* | ACAGTGGACCTGAACTTCGC | TCGCTGTTCTCCTGCCATTT |
| *Pp1r15a* | GAGAAGACCAAGGGACGTGG | AGCGAAGTGTACCTTCCGAG |
| *Lgals3* | CCCAACGCAAACAGGATTGT | AGTTGGCTGATTTCCCGGAG |
| *Slc7a11* | TGCATATGCTGGCTGGTTTT | CTCCGCACTGATGGTGGTAA |
| *Anxa1* | CCAGCAGGAGCTTTCCTCAT | GAGTTCCGGCACCCTTCAT |
| *Msra* | GGGGTGACGTCTGTAGGAAG | GTTTTCTTGCAGCCCCACAG |
| *Egr1* | GAGCACCTGACCACAGAGTC | AAAGGGGTTCAGGCCACAA |
| *Atf3* | AGTGCCTGCAGAAAGAGTCA | GTTCCTCTCGTCTTCCGGTG |
| *Hmox1* | CAGAGCCGTCTCGAGCATAG | AAATCCTGGGGCATGCTGTC |
| *Gclm* | CAATGACCCGAAAGAACTGCT | GACAACAGCAGGTCGGTGA |
| *Txn1* | TGTGGATGACTGCCAGGATG | TCCTTGTTAGCACCGGAGAA |
| *Srxn1* | ACACGATCCTGGCGGACC | CTCACGAGCTTGGCAGGAAT |
| *Alox12* | TGGCTAAGATCTGGGTCCGA | GTGTGGAACGAGGAGCTTGA |
| *Bst2* | GTCACGAAGCTGAACCAGGA | CAAACAACTGTGACCTGCCA |
| *Dhx58* | GAACCCCAACTTCTCGGTCT | TCCGATTTTGAGCACTGGCA |
| *Oas1g* | CTCCAAGGTGGTGAAGGGTG | CTTGAAGCTCAGAGACCGGG |
| *Gas6* | GGCTCAACTACACCCGAACA | CCAGGGCAACATCCTCAACT |
| *Sfrp1* | CTGGCCCGAGATGCTCAAAT | ACGGTTGTACCTTGGGGC |
| *Igf1* | CCAAGACTCAGAAGTCCCCG | GTGGCATTTTCTGCTCCGTG |
| *Oas2* | TCTGAAGCAGATTGCGGTGT | ATAGGAGCCACCCTTAGCCA |
| *Bmp4* | GGTCTCCGTCCCTGATGGG | CCAGGAATCATGGTGTCTTG |
| *Stat2* | TCCGCTGTTCGCTATCTTGG | TGCGCCATTTGGACTCTTCT |
| *Irgm2* | CTTCTGAGCAGCCACCCC | CCCTTCTTTCACGGCAGTCT |
| *Stat1* | CGCTGCCTATGATGTCTCGT | TTCCCTCCTGGGCCTGATTA |
| *Il12rb1* | ACCATGCAAGGACAGTCACC | TGGATGTCATGTTGCCTCCC |
| *Irf1* | GGGACATAACTCCAGCACTGT | GGCATGGGTGACACCTGTAG |
| *Oasl1* | CAGGGGACAAAGGCCTATCA | CACGGTCACCTGGATATCGG |
| *Irf8* | GCGTCTGTCCAACTGCTTTG | CGGTCACACATCCTGCAATC |
| *Oasl2* | GCTTACGATGCTTTGGGACC | CCAACCGGAGGAGGTTCTTC |
| *Jak1* | TTTAGTCCCATGGCCTTGTTCC | AATGGCTTGGGAGAGAAGGA |
| *Stat3* | CAGGAGGGCAGTTTGAGTCG | CAGAGGTGCTCTCCTCTTTAC |
| *Nkg7* | CATGGCTTTTTCTGCAGCTCTC | AGGACAGGCTCCAAACATCC |
| *Prf1* | GGTGGGACTTCAGCTTTCCA | TGCTTGCATTCTGACCGAGT |
| *Il12rb2* | CCAGGGAGCATCACGAAGTT | CACTCACTGGTGCTTTGTGC |
| *Gzmb* | GGACATGAAGTCAAGCCCCA | TCTTGGCCTTACTCTTCAGCT |
| *Gzma* | AAAGGACTCCTGCAATGGGG | ATCGGCGATCTCCACACTTC |
| *Nf-κB* | GAAGATGAGGGAGTGGTGCC | GCTTCATGTCCCCTTGTGGT |
| *CD8A* | CATCTGCTACCACAGGAGCC | CTGGCGGTGCCATTTTACAC |
| *Caspase8* | CCAGATTTCTCCCTACAGGGT | TTTTCTGCCAGCATGGTCCT |

**Supplementary Figures**

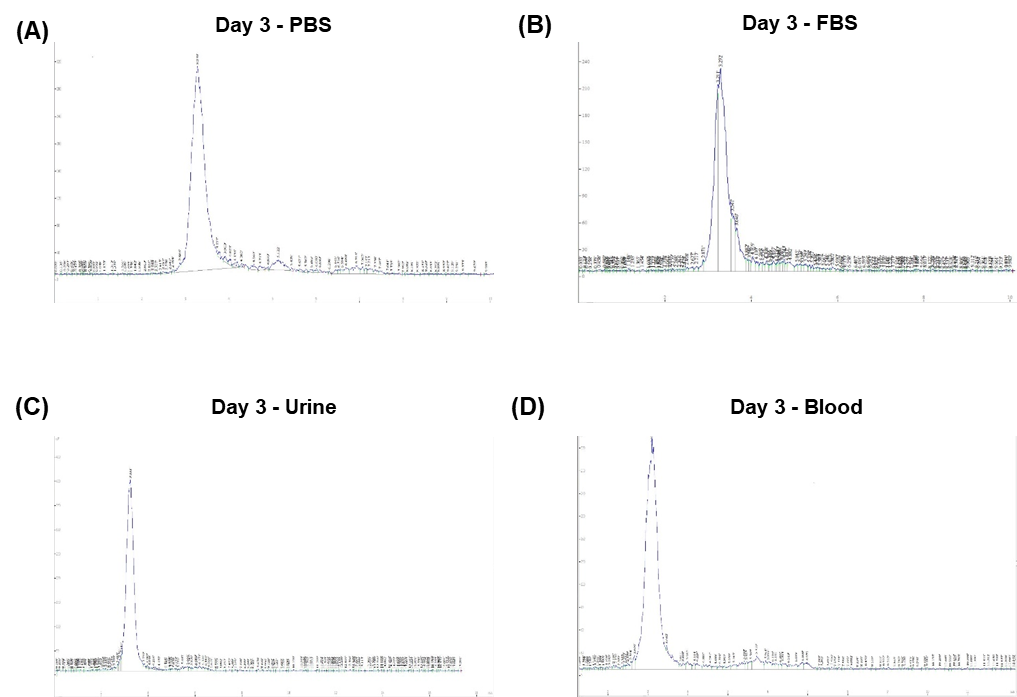

**Supplementary Figure S1.** The stability assessment of ²²⁵Ac-FAPI-04. (A) *In vivo* stability of ²²⁵Ac-FAPI-04 in PBS after injection 72 h after injection by Radio-TLC. (B) *In vivo* stability of ²²⁵Ac-FAPI-04 in FBS after injection 72 h by Radio-TLC. (C) *In vivo* stability of ²²⁵Ac-FAPI-04 in urine after injection 72 h after injection by Radio-TLC. (D) *In vivo* stability of ²²⁵Ac-FAPI-04 in blood after injection 72 h after injection by Radio-TLC.

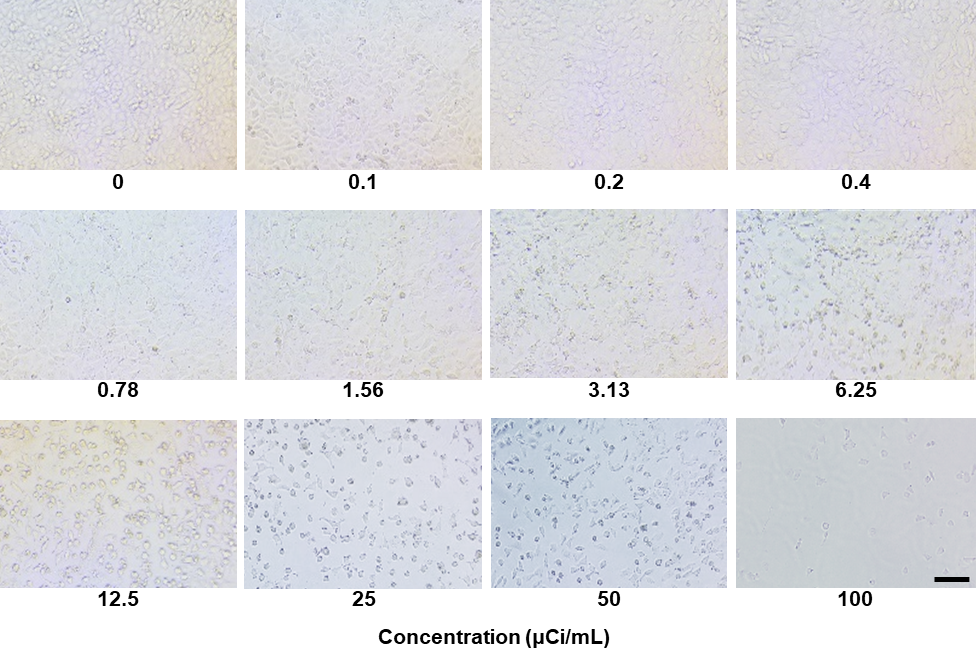

**Supplementary Figure S2.** Imaging of the half-maximal inhibitory concentration of ^225^Ac-FAPI-04 in B16F10. scale bar, 100 μm.

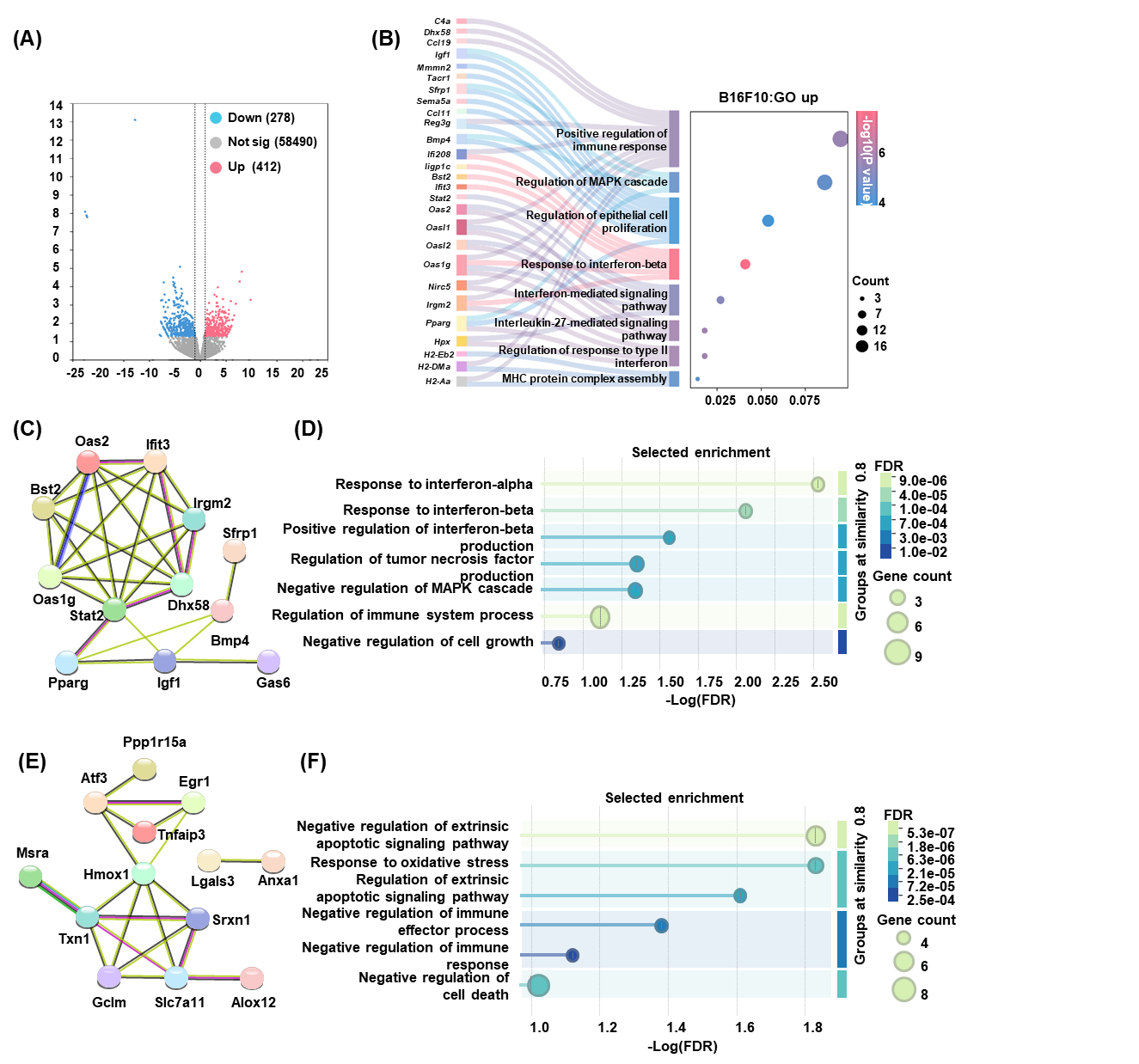

**Supplementary Figure S3.** Transcriptomic analysis of regulated genes and their protein-protein interaction network. (A) Volcano plot of DEGs between the control group and the treatment group. The cutoffs used in the analysis are FDR < 0.05 and |log2 fold change| > 1. (B) The Sankey diagram showed genes that are upregulated in enrichment GO pathways related to immune response, MAPK, and interferon. The cutoffs used in the analysis are FDR < 0.05 and q-value < 0.05. (C) The protein-protein interaction network regulated by the upregulated genes showed the interactions between proteins. Confidence score > 0.7 and FDR of GO pathway < 0.05. Created in STRING. (D) The GO enrichment of upregulated genes in Fig.S3C related to the immune and interferon pathway. The cutoffs used in the analysis are FDR < 0.05 and q-value < 0.05. (E) The protein-protein interaction network regulated by the downregulated genes in Fig.S3E showed the interactions between proteins. Confidence score > 0.7 and FDR of GO pathway < 0.05. Created in STRING. (F) The GO enrichment of downregulated genes related to the immune and apoptosis pathways. The cutoffs used in the analysis are FDR < 0.05 and q-value < 0.05.
